# THE ROLE OF LIQUID CRYSTAL ORDERING IN THE STRUCTURAL ORGANIZATION OF DNA IN BACTERIA

**DOI:** 10.64898/2026.08.31.748243

**Authors:** Yu. F. Krupyanskii, V. V. Kovalenko, N. G. Loiko, A. A. Generalova, E. V. Tereshkin, K. B. Tereshkina, O. S. Sokolova, G. S. Peters

## Abstract

This paper presents and critically reviews the results of original and some literature based experimental studies conducted by the authors last years on the structural organization of DNA in dormant (starvation stress), anabiotic dormant (4 HR treatment) E. coli cells, as well as the K12 &[Delta]dps strain, which lacks the Dps protein (Dps null E. coli). The experimental data includes small-angle synchrotron radiation diffraction (SAXS) and transmission electron microscopy (TEM) data. Synchrotron radiation diffraction experiments on K12&[Delta]dps cells allowed us to conclude that peaks at 44.3, 22.1, and 14.8 angstrom resolutions are associated exclusively with ordered DNA organization. Peaks at 44.3, 22.1, and 14.8 angstrom resolutions are also observed for samples of dormant (starvation stress) cells and anabiotically dormant cells. Therefore, this ordered DNA organization also applies to samples of dormant and anabiotically dormant cells. A model is proposed that considers the ordered DNA organization in the cell as a cholesteric liquid crystal. The powder diffraction pattern calculated based on this model is compared with experimental small angle X ray scattering (SAXS) data obtained on Dps-null cell samples. The model completely reproduces the key features of the experimental diffraction pattern from Dps-null cell samples. Accordingly, the cholesteric liquid crystal model corresponds to DNA packaging in dormant and anabiotically dormant cells. Cholesteric liquid crystal ordering should be further considered in all models of cellular DNA packaging. To address the question of which structural organization of DNA predominates in the cell: the cholesteric liquid crystal or nanocrystalline or whether they coexist and fully manifest themselves under different external conditions, it is necessary to utilize the latest methodological advances in structural analysis.

## I. INTRODUCTION

3D genome architecture determines cell function. Studying the mechanisms of changes in DNA structural organization in bacteria in response to various types of stress allows us to understand not only the fundamental mechanisms of DNA structural organization (3D architecture) in cells, but also the mechanisms of bacterial survival, and to advance the solution of an important medical problem: overcoming the resistance of pathogenic bacteria to drugs (including antibiotics) [1].

Bacterial genomic DNA interacts with nucleoid-associated proteins (NAPs) and is *organized* in a highly condensed and functional form in the cell’s nucleoid. Five decades of intensive research have shown [2] that DNA is *organized* hierarchically in the nucleoid of actively growing Escherichia coli (E. coli) cells, with three levels of compaction [2, and references therein].

This *organization,* or high level of dynamic order, is characteristic of actively growing cells, which can maintain this state through metabolism [3, 4]. Since this *organization* (high level of *dynamic order*) is critically dependent on constant energy consumption, it cannot be maintained in stressed, and therefore energy-depleted, bacteria in a stationary state. This article examines the structural *organization* of DNA in cells exposed to starvation stress and a chemical analogue of the autoinducer of anabiosis (4-hexylresorcinol, 4-HR). Under severe stress, energy resources are insufficient for normal biochemical protection of DNA, so cells are forced to use another energy-independent mechanism for maintaining order to protect DNA (like inanimate nature). Under certain concentrations and conditions, DNA macromolecules (in complex with Dps (DNA-binding protein from starved cells) or on their own) tend to self-organize into ordered structures— liquid crystalline structures, crystals, etc. [4]).

The structure and properties of DNA in an actively growing cell are, to a first approximation, well modeled by a folded (or fractal) globule, characterized by a hierarchy of folds and forming a self-similar structure [5,6].

For dormant cells, a completely new structural organization of condensed DNA should be expected compared to the organization in actively growing cells. In almost all the works we cite [4,7–10, 12–16], the priority mechanism of DNA protection under stress is still considered to be “biocrystallization” or nanocrystallization of DNA with the Dps protein [4,16]. The present work is devoted to a critical analysis of the existing models of DNA packaging in the cell (under starvation stress and the action of 4-hexylresorcinol, 4-HR). For this purpose, extensive original and literature experimental material on synchrotron radiation diffraction [7–11] and numerous data on electron microscopy [4,9,12, 14–16] will be used. For a better understanding of the below-mentioned in this article and the resulting cholesteric liquid crystal model of the structural organization of DNA in the cell, the work provides brief information on the process of bacterial growth, the Dps protein, the change in the ordering model and the formation of dormant cells. A short review of the cholesteric liquid crystalline ordering of DNA is given. Next, we will discuss the key experimental data necessary for a critical analysis of existing models, and formulate a new cholesteric liquid crystal model of DNA structural ordering. The role of cholesteric liquid crystal ordering in the structural organization of DNA in cells will be elucidated.

### II.1 BACTERIAL GROWTH. Dps PROTEIN

Bacterial growth is the division of a bacterial cell into two daughter cells. The daughter cells are genetically identical to the parent cells. The dynamics of bacterial population growth are divided into four phases [17]. The first growth phase is called the lag phase, followed by the exponential phase, during which rapid exponential population growth occurs. During the exponential phase, nutrients are consumed at maximum rate until one of the essential compounds is depleted and begins to suppress growth. The third growth phase, called the stationary phase, begins when nutrients for rapid growth are insufficient. The metabolic rate drops, and cells begin to break down proteins that are not strictly essential. In this phase, cells begin to experience starvation stress. In response to starvation stress, microbial cells activate hereditary adaptation strategies that allow them to preserve part of their population and survive under any adverse conditions. These strategies are aimed at protecting the cell’s genetic material (DNA) [18]. If starvation stress persists, either dormant forms, characterized by a virtually complete absence of metabolism (exchange with the external environment), or cell death occur. Most cells (up to 99.98%) in long-starved populations die. The remaining cells (0.02%) develop into dormant forms [1, 12]. In the exponential growth phase, the relative content of DNA-associated histone-like proteins (NAP) begins to change. While the Dps (DNA-binding protein of starved cells) protein content during the growth phase is approximately 6,000 proteins per cell, in the stationary and late stationary phases, the Dps content becomes overwhelming compared to other proteins—180,000–200,000 proteins per cell. Dps plays a regulatory and protective role in E. coli cells [19–22]. During starvation, Dps is highly active and can significantly alter the structure of bacterial DNA.

### II.2. THE CHANGE OF THE ORDERING MODEL. FORMATION OF DORMANT CELLS

Living systems maintain an ordered, far-from-equilibrium state by feeding on chemical compounds. Metabolism (exchange with the external environment) is the source of *dynamic* order. If metabolism ceases, thermodynamic equilibrium cannot be avoided. As cells transition to a dormant state (a virtually complete absence of metabolism), the usual biochemical mechanisms for DNA protection cease to function. Cells, adapting to new conditions, are forced to use physical mechanisms to protect DNA (dense DNA packing (cholesteric liquid crystalline organization), nanocrystallization of DNA with proteins, etc.). These adaptive mechanisms are necessary for cell survival. When favorable conditions return, surviving cells will resume normal life. Dormant cells can be expected to have a completely new structural organization of DNA compared to growing cells.

## III. MATERIALS AND METHODS

### III.1. Studied samples

Below is a brief description of the samples for which the experimental results are presented in this article. The following E. coli bacterial strains were used: 1 – K-12MG 1655, wild-type strain; 2 – K-12 MG1655 Δdps, Dps-null mutant; 3 – Top10, wild-type strain; 4 –Top10/pBAD-DPS, a mutant with overexpression of the Dps protein; 5 – BL21-Gold(DE3)/pET-DPS, a mutant with overexpression of the Dps protein.

The reagents used and the procedure for obtaining genetically modified E. coli strains, as well as the preparation of dormant E. coli cells for various strains, are described in detail in [7, 12, 14]. The process of accumulation of the recombinant DPS protein is described in detail in [7, 12].

4-Hexylresorcinol (4HR), chemical formula 4-hexylbenzene-1,3-diol (Sigma-Aldrich, St. Louis, MO), was used as a chemical analogue of the anabiosis factor. The procedure for culturing bacteria, obtaining anabiotic and mummified E. coli BL21 cells, culturing the anabiotic cells, and preparing bacterial samples for experimental studies is described in detail in [10].

### III.2. Experimental methods

#### III.2.1. Synchrotron Radiation Diffraction

The experiments were performed at station ID23-1 of the ESRF synchrotron (Grenoble, France). The X-ray beam had a wavelength of λ = 1.6799 Å, an aperture width of 10 μm, and an exposure time of 5 s for each diffraction pattern. A PILATUS 6M flat-panel detector was positioned 95 cm behind the sample. The measurements were carried out at a temperature of 100 K.

Additional experiments with the Dps-zero mutant (K-12 MG1655 Δdps) were performed at the BioSAXS (BioMUR) station of the Kurchatov Synchrotron Radiation Light Source, Moscow.

The angle between the axis formed by the incident beam and the diffraction ring is called the scattering angle and is denoted as **2θ**. The scattering vector **q** is given by the formula:

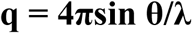

The resolution corresponding to the scattering angle is denoted as **d** and is defined as

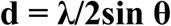

For more details on the method, see [13]

#### III.2.2. Transmission Electron Microscopy

Ultrathin sections were examined using JEM1011 and JEM-2100 transmission electron microscopes (Jeol, Japan) at accelerating voltages of 80 and 200 kV, respectively, and a magnification of 13,000–21,000×. More details on the use of specialized electron microscopy methods for studying DNA-protein complexes in E. coli cells can be found in [14].

## IV. EXPERIMENTAL RESULTS

The experimental results discussed in this article are presented below.

### IV.1. Structural Response to Starvation Stress

#### IV.1.1 Synchrotron Radiation Diffraction

Figure 1 shows the dependences of the synchrotron radiation scattering intensity on samples containing E. coli bacterial cells of the BL21-Gold (DE3) strain transformed with the pET-Dps plasmid and subjected to the induction of increased expression of the Dps protein [7, 8] on the scattering vector **q** using averaging of the 2D diffraction patterns over the azimuthal angle. A sharp and intense peak was found at a characteristic distance with a resolution **d** of approximately 44.5 Å, zones of increased intensity were found at characteristic distances with a resolution **d** of approximately 22.2 and 14.8 Å, and a broad intense peak at a resolution in the region of **d** ∼ 90–93 Å. In the control sample of growing cells, no areas of increased intensity were observed (for more details, see [7, 8]). The results of the diffraction experiments shown in Fig. 1 indicate the presence of an ordered organization of DNA in the bacterial cell with the above-mentioned values of resolution **d**.

**Figure 1.**
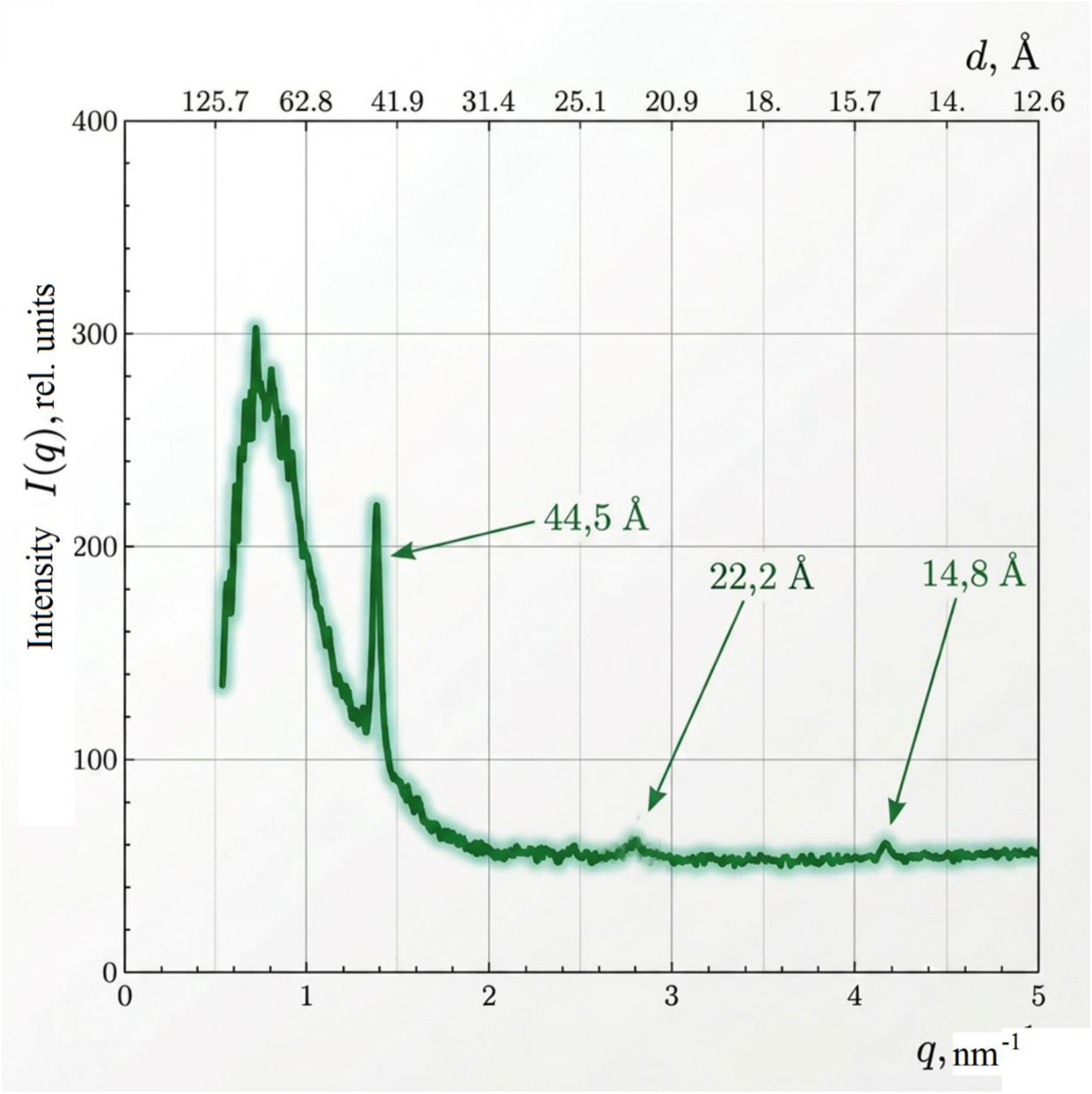
Scattering intensity as a function of the scattering vector q for a sample of starved E. coli strain BL21-Gold(DE3).

#### IV.1.2. Transmission Electron Microscopy

Figure 2a shows a section of a tomogram of a cell with an ordered structure. The inset in Figure 2a shows the result of a Fourier analysis of the ordered region of the cell, outlined by the white border. The result of the Fourier analysis most likely indicates the presence of a nanocrystal in this region of the cell [12]. A filtered DNA-Dps crystal is shown in Figure 2b. The inset to this figure shows the intensity profile of the electron density along the white line of the main image. The electron densities, apparently corresponding to the interlayer DNA strands, are highlighted in black [12]. The authors of [16] hypothesized that DNA is localized between hexagonally packed Dps layers in the crystal. This means that the characteristic distance between DNA-DNA strands is approximately 90 Å. We note a detail important for the subsequent discussion: DNA is not directly visible in electron microscopic studies, therefore the proposed information about the location and conformation of DNA in the cell is hypothetical.

**Figure 2.**
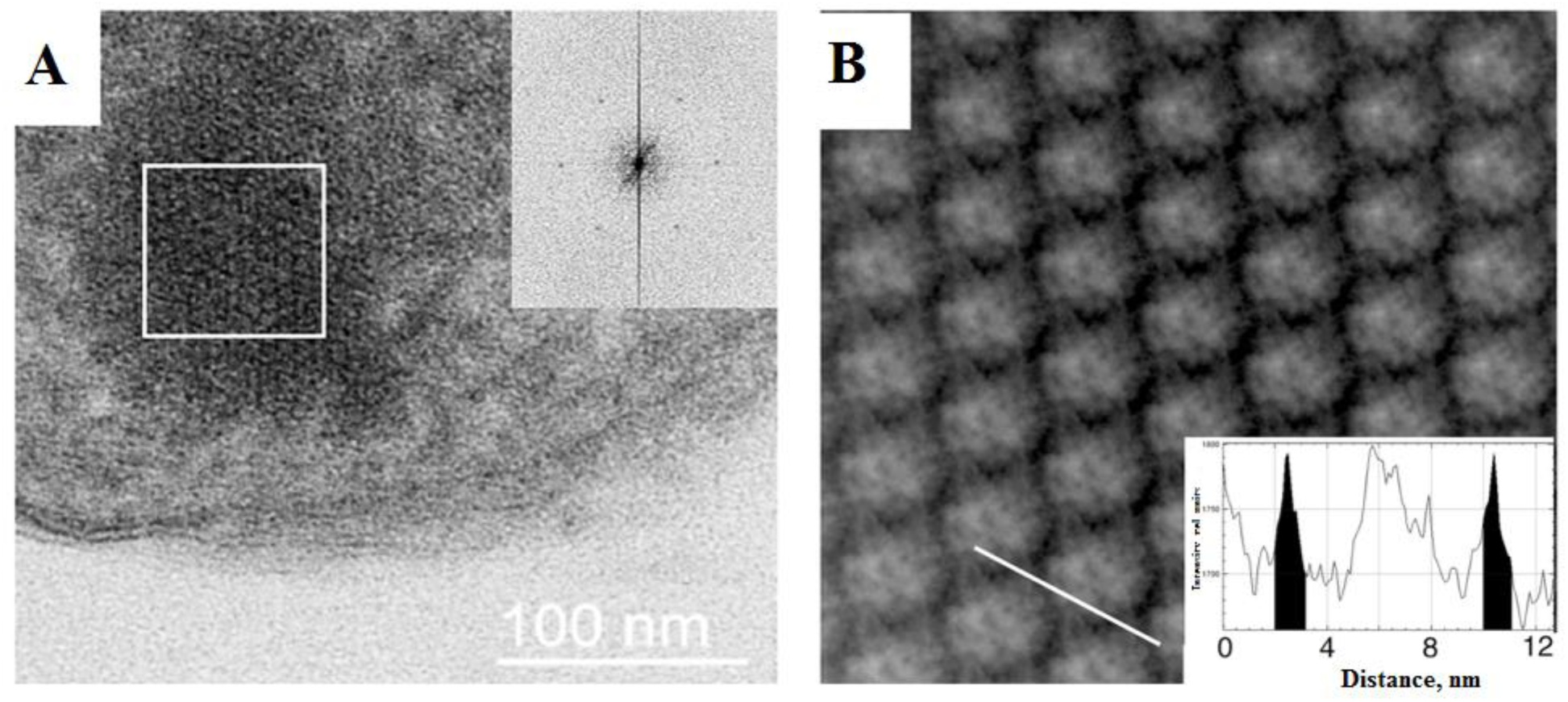
a – Cell tomograms (the inset shows the result of Fourier analysis of the cell region outlined by the white border); b – filtered DNA–Dps crystal.

#### IV.1.3. Alternative Types of DNA Condensation in Dormant Cells

Using transmission electron microscopy and dual-axis tomography, three types of novel (compared to growing cells) condensed DNA–Dps structures were detected in dormant E. coli cells. These structures include nanocrystalline DNA structures, liquid crystalline DNA structures, and folded nucleosome-like DNA structures (see Fig. 3a, b, and c, respectively). Energy-dispersive spectroscopy (EDS) was used to detect and position the desired elements in selected cellular regions. For example, the Kα peak (2.307 keV) for sulfur reflects the presence of the DNA-binding protein Dps (each Dps protein contains 48 methionines), while the Kα peak (2.013 keV) corresponds to phosphorus in DNA. The simultaneous presence of both peaks in the structure spectra most likely indicates the formation of a DNA– Dps complex. Representative EDS spectra are shown in Fig. 3d. The relative atomic composition of different regions of the samples is proportional to the intensity of the EDS peaks. It is noteworthy that liquid crystalline structures (illustrated in Fig. 4) were detected in dormant E. coli bacterial cells [12] in all their populations: both with the dps gene and without it (Dps null), i.e., in the absence of the Dps protein in the cell. In some cells, the structure of condensed DNA resembles a cholesteric liquid crystal [12]. Dense packing of DNA in the liquid crystalline phase reduces the accessibility of DNA molecules to various damaging factors, including radiation, oxidants, and nucleases [4]. Note that the liquid crystalline type of DNA condensation is the only case (except for DNA toroids [16]) when condensed DNA molecules in a cell can be directly seen in an image. Of particular interest is the third type of ordered structure, first discovered in our study [12] in dormant E. coli cells: a folded nucleosome-like structure (see Fig. 3B). A detailed description of this folded nucleosome-like structure is beyond the scope of this article and can be found in [12, 13]. What is important here is that the synchrotron radiation diffraction pattern from this structure is most likely similar to the diffraction pattern from a growing cell, i.e., it does not exhibit any distinctive features (diffraction peaks).

**Figure 3.**
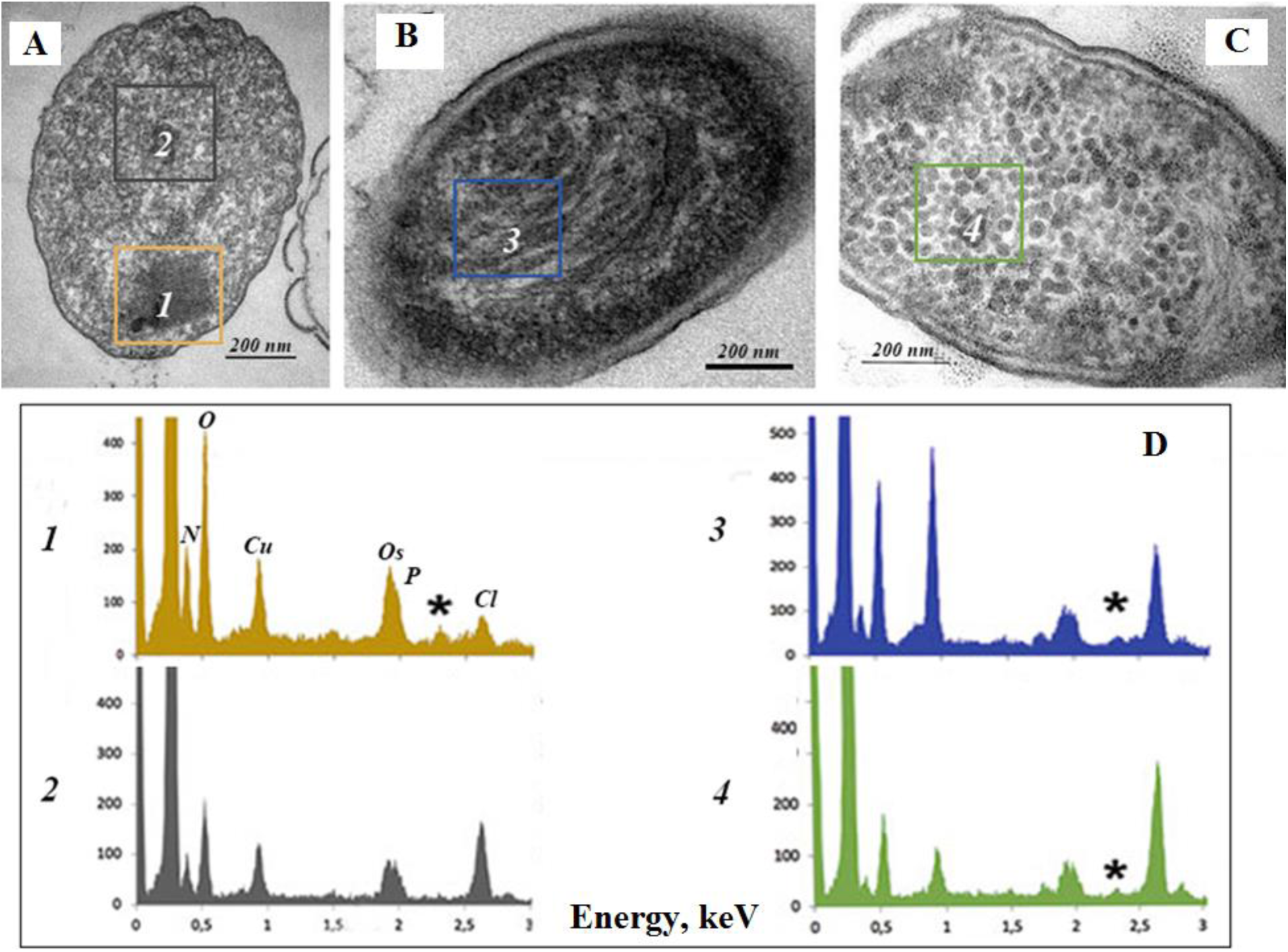
Three types of condensed DNA–Dps structures detected in dormant E. coli cells (starvation stress): a – nanocrystalline; b – liquid crystalline; c – folded nucleosome-like type; d – EDS spectra from selected regions (marked with colored frames in Figs. a–c). The positions of the S peak are marked with an asterisk.

**Figure 4.**
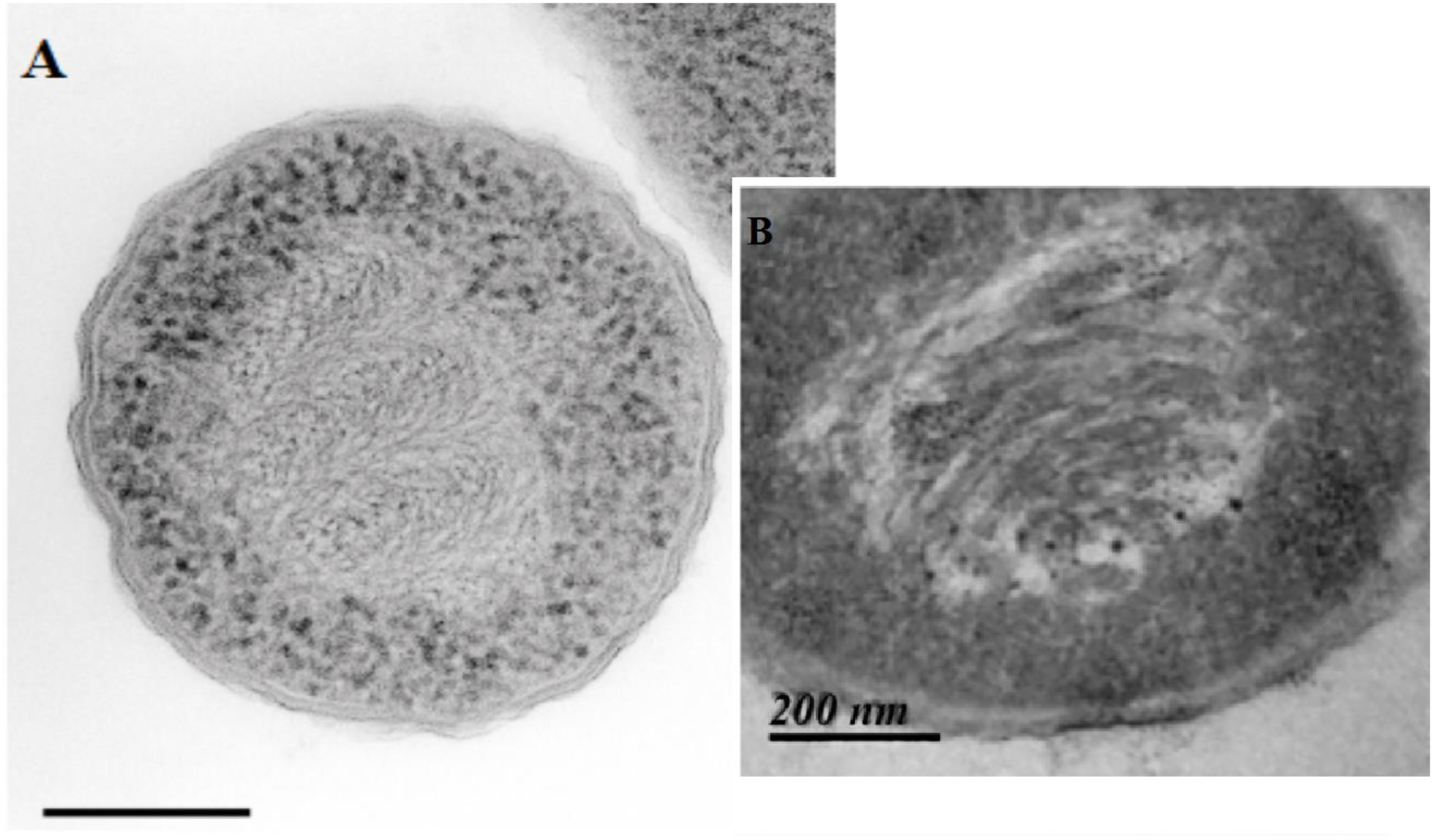
DNA–Dps liquid crystalline assemblies in dormant E. coli cells: A – starved E. coli cells (Dps-null mutant, Dps proteins are absent), the cholesteric liquid crystalline phase of DNA is visible: DNA (in the form of nested arcs characteristic of the cholesteric phase) and ribosomes, which appear as dark particles at the periphery of the cell (taken, with permission, from [4]); B – Top10/pBAD-Dps strain (M9 medium, induced Dps production in the linear growth phase [12]);

Table 1 reflects the tendency (as a percentage of the total number of cells) for the formation of a particular condensed DNA structure for cells of a particular strain and growth conditions after prolonged starvation. We would like to emphasize an important observation for further study: the liquid crystalline structures shown in Fig. 4, were detected in dormant E. coli bacterial cells [12] in all populations: both with and without the dps gene (Dps null), i.e., in the absence of the Dps protein in the cell.

**Table 1.**
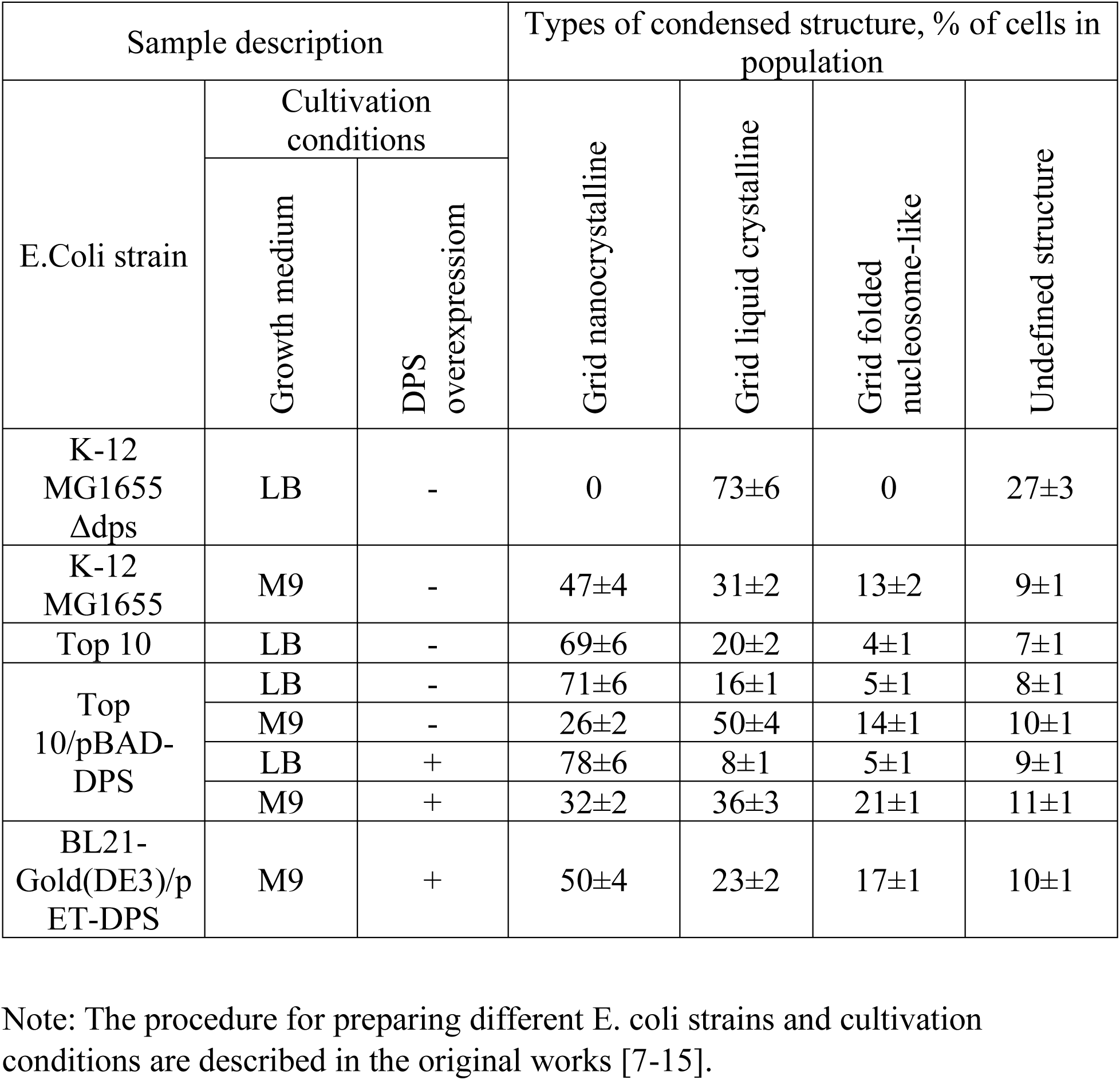
Different types of DNA structural organization in the studied dormant E. coli cells.

#### IV.1.4. Structural organization of DNA upon exposure to a chemical analogue of the anabiosis autoinducer, 4-hexylresorcinol (4-HR)

We will consider the structural organization of condensed DNA in cells exposed to a different type of stress than starvation: exposure to a chemical analogue of the anabiosis autoinducer, 4-hexylresorcinol (4-HR). Several studies have shown that the introduction of 4-HR at concentrations of up to 10^−4^ M into a culture of growing cells leads to a cessation of cell division and their premature transition to stationary phase [10]. A further increase in the 4-HR concentration in the cell culture initiates the transition of cells into a dormant anabiotic state (almost complete absence of metabolism), reminiscent of the dormant state of cells under starvation stress. Anabiosis is understood as the suspension of cellular activity followed by its restoration under favorable conditions [10]. A further increase in the 4-HR concentration causes a complete loss of viability of bacterial cells. These cells are called micromummies [23]. The preparation of anabiotic dormant cells is described in detail in [10].

##### IV.1.4.1. Synchrotron Radiation Diffraction Data

Figure 5 shows the synchrotron radiation scattering curves for cells exposed to 4-HR at a concentration of 10^−4^ M (blue curve). The scattering curve obtained from the growing cell population, in our case taken as the reference (control) curve against which all observed diffraction effects were considered, is shown in the same Figure 5 in yellow. The scattering intensity curve from E. coli cells in the stationary growth phase exposed to 4-HR at a concentration of 10^−4^ M is identical in shape to the curve obtained from cells under starvation stress (Fig. 1). The curve shows peaks corresponding to d resolutions of 44, 22, and 14.7 Å, and a broad peak with a maximum intensity at a resolution of ∼90 Å. This coincidence of the diffraction curves suggests that the structure of condensed DNA within anabiotic and dormant (starvation stress) cells is identical, and that the mechanism of formation of protective ordered structures is very similar.

**Figure 5.**
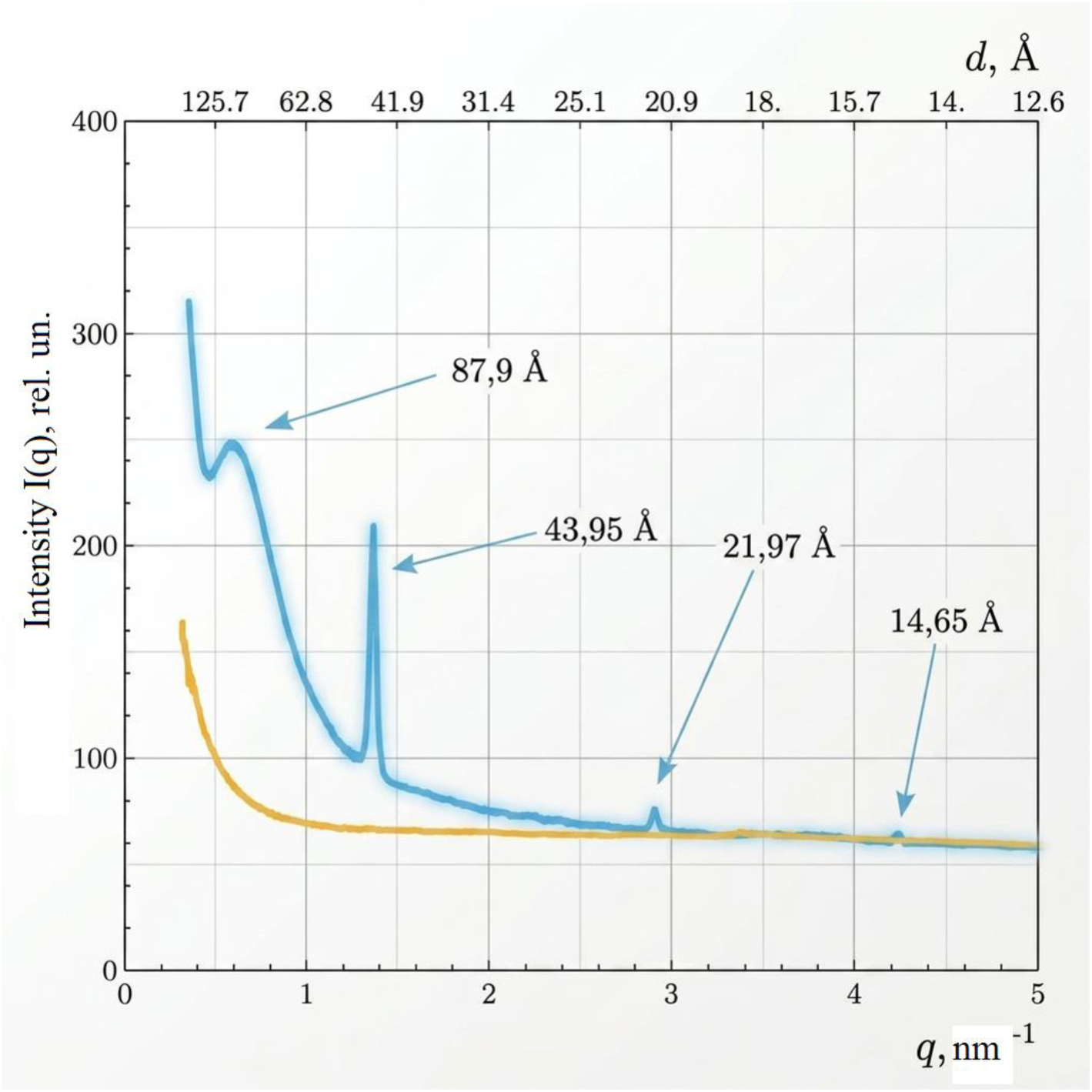
Scattering intensity as a function of q for samples containing E. coli Gold bacterial cells exposed to 4-HR at a concentration of 10–4 M during the stationary growth phase, anabiotic state of the cell (blue curve); The yellow curve (in this figure) is the scattering intensity from cells in the active growth phase.

##### IV.1.4.2. Transmission Electron Microscopy Data

Figure 6 shows electron micrographs of E. coli cell sections (Top10/pBAD-DPS) in the stationary growth phase after the addition of 4-HR at a concentration of 10^−4^ M: A – without Dps protein induction, B – with Dps protein induction. The inset in Fig. 6B shows the Fourier transform of the highlighted region of the cell. Note two characteristic features of DNA packaging in the images shown in Fig. 6A and 6B. Firstly, these images clearly show DNA; secondly, the DNA packaging has a characteristic arcuate shape, which is typical of cholesteric liquid crystalline DNA packaging (compare with the DNA packaging shown in Fig. 4).

**Figure 6.**
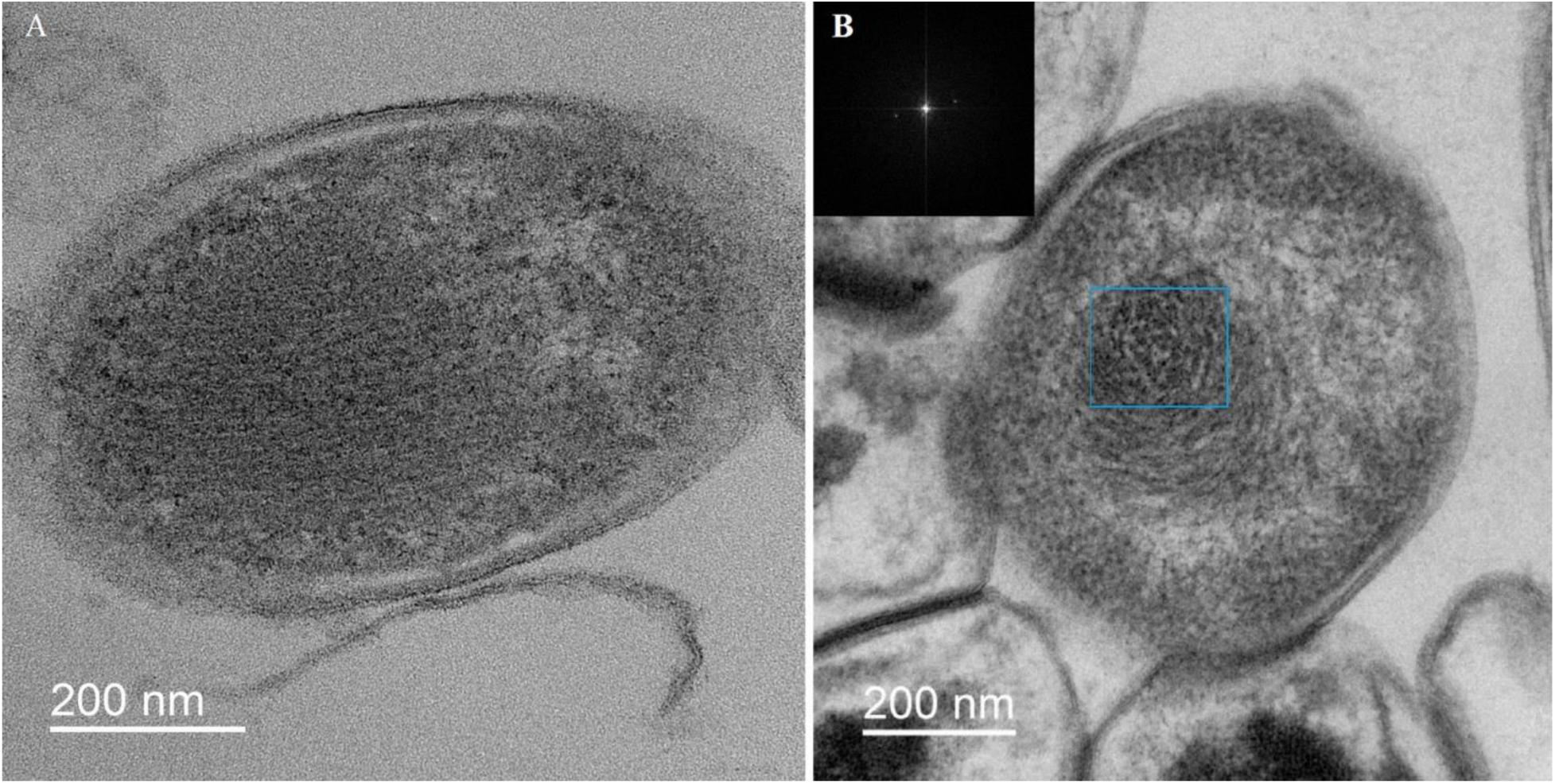
Electron micrographs of E. coli Top10/pBAD-DPS cell sections during the stationary growth phase in cultures: A – without Dps protein induction, B – with Dps protein induction. The inset shows the Fourier transform of the highlighted region. Fourier analysis indicates that the structure in the highlighted region is not crystalline.

Original experimental studies allow us to conclude that: 1) 4-HR initiates the cell’s transition to an anabiotic dormant state; 2) the dormant anabiotic state is structurally very similar to the dormant state obtained as a result of starvation; 3) In the dormant anabiotic state, both cholesteric liquid crystalline DNA structures and nanocrystalline DNA structures are observed. DNA is not visualized for nanocrystalline structures (see Fig. 10 in [24]). Visible nanocrystals are most likely formed by associated proteins.

#### IV.1.5 Structural Organization of DNA in Dps-Null Mutant Cells E. coli K-12 MG1655Δdps (hereinafter K-12 Δdps)

The procedure for constructing and preparing the K-12 Δdps strain is described in detail in our previous studies [12, 11].

##### IV.1.5.1. Synchrotron Radiation Diffraction Data

Let us examine the results of the synchrotron radiation scattering experiment on the E. coli K-12Δdps strain (a strain lacking the Dps protein). Figure 7 shows the dependence of scattering intensity on the scattering vector for this strain in the stationary phase. Dps protein is absent from the cell; however, Figure 7 shows diffraction maxima at 44.3, 22.1, and 14.8 Å, i.e., in the same positions as in Fig. 1 and Fig. 5. This means that the Dps protein is not associated with the formation of the diffraction peaks at 44.3, 22.1, and 14.8 Å. Consequently, these peaks are associated only with the ordered organization of DNA and are of greatest interest for understanding DNA packaging in the cell.

**Figure 7.**
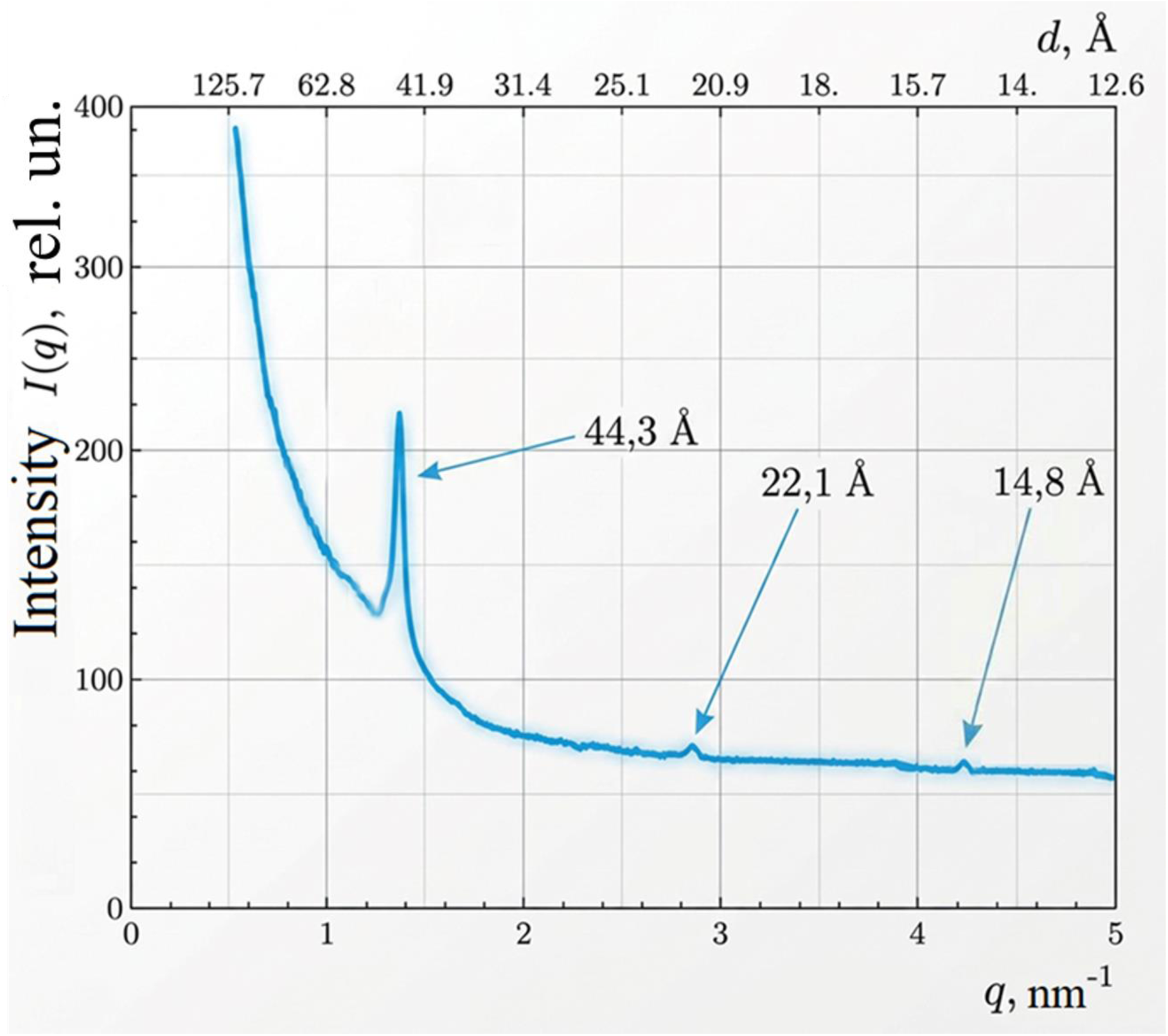
Scattering intensity versus scattering vector q for a sample of the K-12 Δdps strain (Dps-null, no Dps proteins)

## V. RESULTS

### V.1. Critical analysis of the results on the structural organization of DNA in the cell

Let us turn to models of DNA packaging in E. coli bacterial cells during the stationary phase. In this phase, the main DNA-binding protein is Dps, the number of which approaches 200,000 per cell. The interaction between Dps and DNA leads to the rapid formation of highly ordered, densely packed DNA-Dps associates. Figure 8a reflects the first model for the formation of DNA-Dps nanocrystals on a toroidal DNA structure serving as a template for further DNA-Dps crystal growth [4,16].

**Figure 8.**
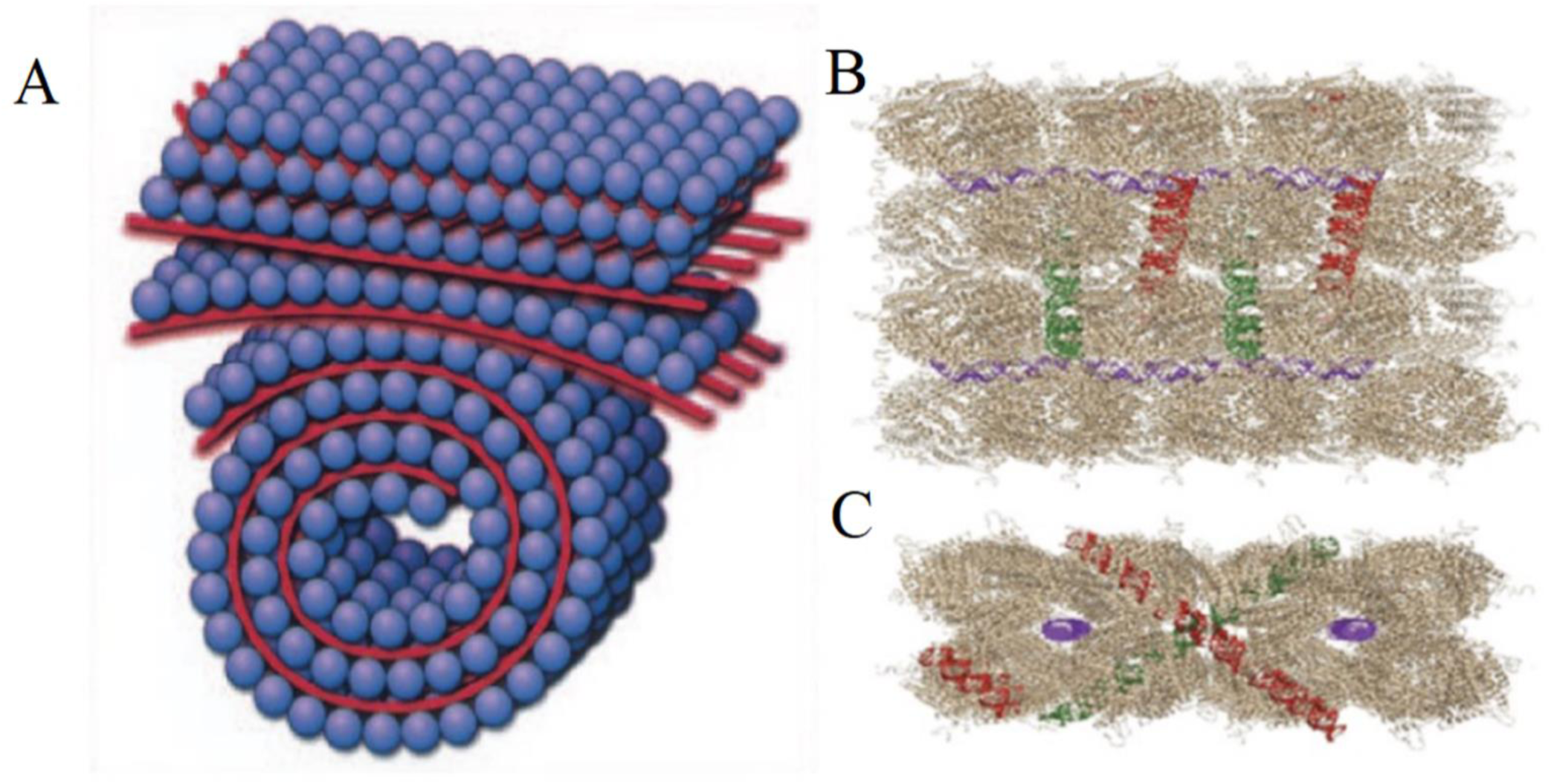
A - Model of the intracellular DNA–Dps assembly depicts the initially formed toroidal structure, which acts as a template for the DNA–Dps crystal (adopted with permission from [4]). Possible DNA–Dps arrangements: B - top view, C - side view.

DNA (red bands) is localized between dodecameric Dps particles (blue spheres with a diameter of 90 Å). This means that the distance between the DNA strands is approximately 90 Å. The diffraction pattern of DNA molecules located relative to each other at 90 Å should consist of a main sharp (narrow and intense) peak with a resolution of 90 Å and should have a noticeable satellite corresponding to a resolution of 30 Å. Nothing similar was observed in any of the experiments presented. Thus, the first model does not satisfy the results of the diffraction experiments.

Possible arrangements of DNA relative to Dps are shown in Fig. 8B [13,15]. It should be noted that this second model almost completely repeats the ideas of the first model given in [16] (Fig. 8A), therefore all the shortcomings of the first model are also inherent in the second model and its modifications [25].

To clearly illustrate the indicated shortcomings of the first and second models, Fig. 9 shows the calculated diffraction curve of the scattering intensity as a function of the scattering vector **q** for a system consisting of 16 Dps proteins and 16 double-stranded (ds) DNA molecules located between them. Figure 9 clearly shows a prominent satellite at 29.8 Å, which was not observed in our experiments. Note that the system used to calculate the diffraction curve for Figure 9 is very small (16 Dps – 16 DNA). It is known that only for relatively large systems [26] do the main peaks (in our case, ∼90, 45, 22, and 15 Å) stand out noticeably from the interpeak beats, whereas for small systems, the beat intensities can be up to 20% of the main peak intensity.

**Figure 9.**
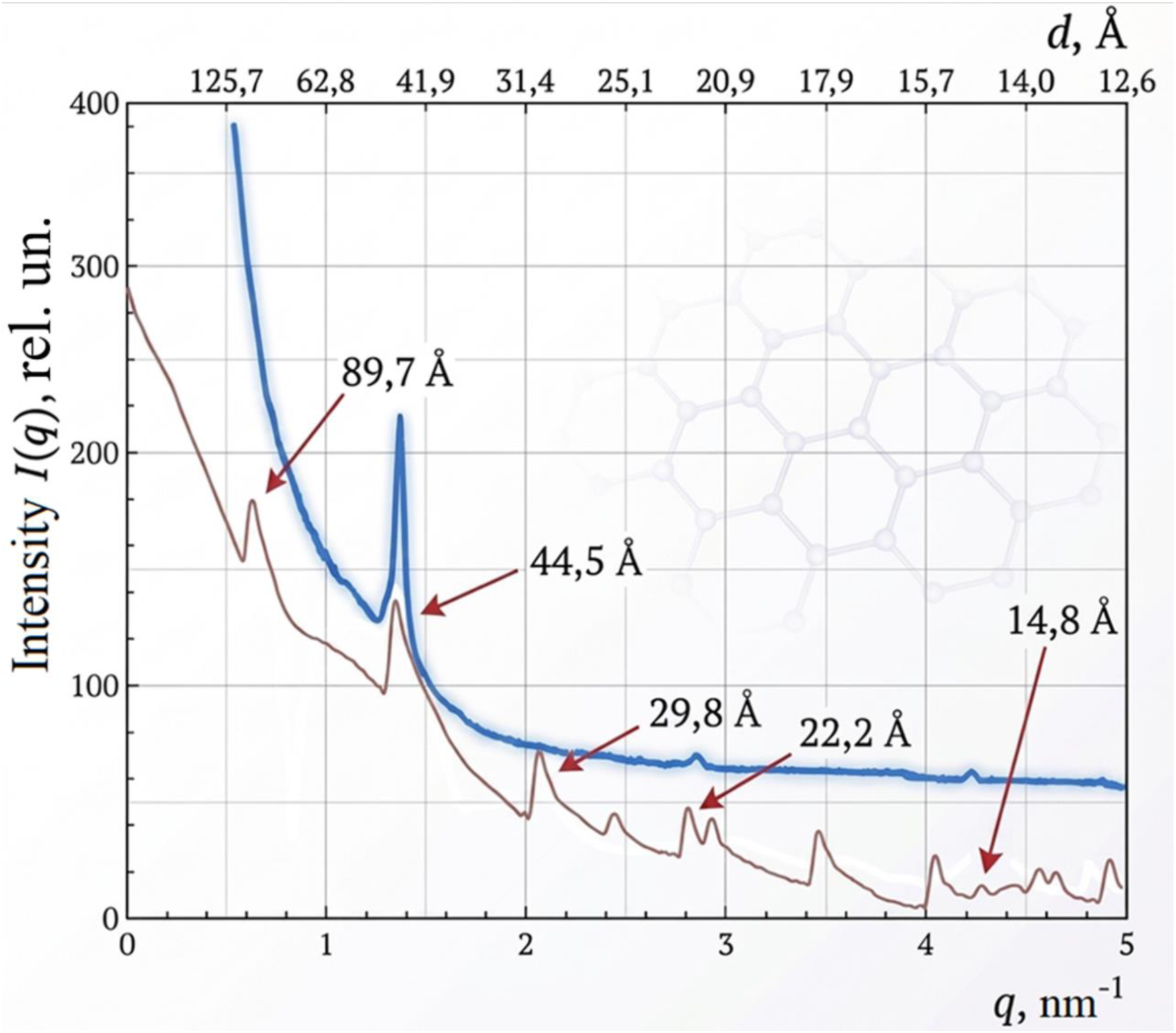
Scattering intensities versus scattering vector q: experimental curve (blue curve) for a sample of the Dps-null strain, and the calculated diffraction curve (red curve, intentionally positioned just below the experimental curve) for a system consisting of 16 Dps proteins and 16 double-stranded (ds) DNA molecules located between them.

### V.2. Model of DNA Structural Organization in Dps-Null Cells. Description of Experimental Data for the K-12 Δdps Strain

Before moving on to the model describing the original data for Dps-null, we present the basic information about the cholesteric liquid crystalline phase of DNA necessary for further understanding.

#### V.2.1. Cholesteric Liquid Crystal Phase of DNA. [27,28]

It is known that upon reaching a certain critical concentration in aqueous-salt solutions (50–600 mg/ml) at room temperature, double stranded (ds) DNA molecules spontaneously transit from a disordered state to a highly organized, densely packed liquid crystalline structure [29–32]. This process occurs without energy expenditure and is thermodynamically favorable. Such packing, in principle, can be hexagonal. With such dense packing, the existence of DNA “layers” is possible [33, 34], dsDNA molecules lie in the plane of these “layers.” Since the properties of DNA molecules in each hypothetical “layer” are similar to those of nematic liquid crystals, this served as the basis for using the term “quasinematic layer” to designate it [35].

Finally, the presence of several levels of chirality in dsDNA molecules (the helical structure of DNA molecules, the helical arrangement of counterions near these molecules, and the asymmetry of the C-atoms of sugar residues) determines the tendency of neighboring molecules (and, consequently, the “quasinematic layers” of these molecules) to rotate at a small angle relative to each other. Spontaneous rotation of the “layers” of DNA molecules will lead to the formation of a helically twisted (cholesteric) phase, which has properties significantly different from those of the hexagonal phase [36].

The cholesteric phase has three known properties: First, the cholesteric phase arises when the distance between DNA molecules is within 50–20 Å [28], determined using the small-angle X-ray scattering (SAXS) method [31]. Second, a thin layer of the cholesteric phase is characterized by a specific “fingerprint” texture, recorded using a polarizing microscope [37]. Finally, the cholesteric phase exhibits an intense band located in the absorption region of the DNA chromophore (nitrogenous bases), detected using circular dichroism (CD) [38].

The formation of a cholesteric liquid crystalline phase of DNA, arising with increasing DNA molecule concentration, is not described within the framework of standard concepts of liquid crystals of low-molecular compounds. Yu. Evdokimov et al. [28] attempted to evaluate the possible nature of the packing of rigid, linear DNA molecules in the cholesteric liquid crystalline phase. DNA was represented as a rigid rod.

The results of physical and computer modeling of a helical structure consisting of rigid rods fixed in a layer, with adjacent layers rotated by a certain angle, show that sectioning such a structure at a certain angle to the vertical axis of the helix results in the appearance of arc-shaped structures (arches) on the section. These results are schematically depicted in Fig. 10 A, B.

**Figure 10.**
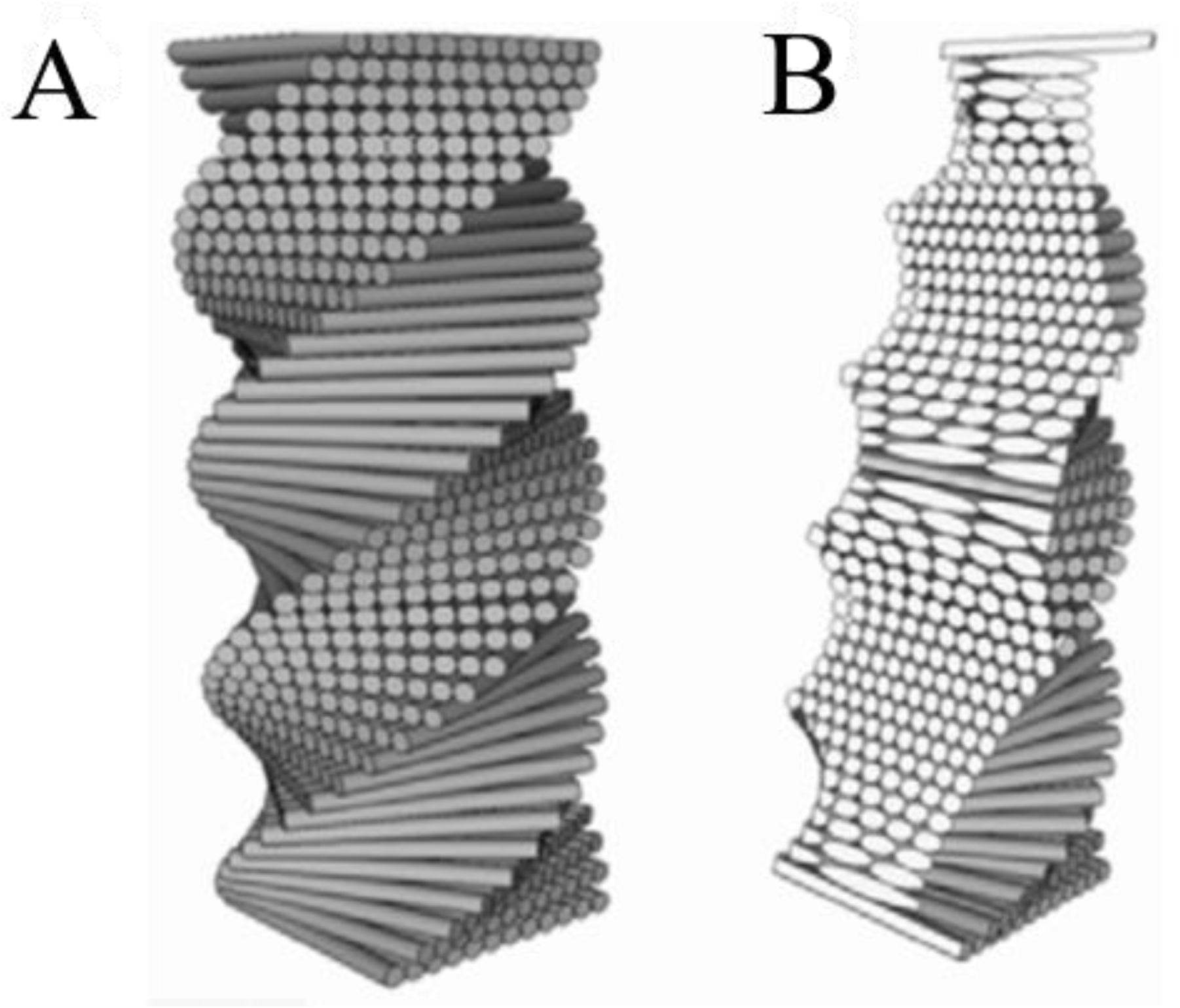
View of a model, spirally twisted structure containing 40 layers of rods, before (A) and after its sectioning at an angle to the vertical axis (B). In Fig. B, a system of equally spaced repeating arcs (“arches”) formed by sections of the rods in the section plane is visible (This figure is a reconstructed figure 7 a, b of the work [28])

There are “biological clues” that indicate that a model like the one presented here is valid. First, there is an electron microscopic study of ultrathin sections of flagellate chromosomes (Dinoflagellata) at different angles of section. These studies revealed molecular patterns, the reconstruction of which unambiguously reflects the cholesteric packing of DNA molecules in this biological object (the patterns form characteristic arches; drawings are provided in the articles [39, 40]). Similar molecular patterns were also observed in sections of chromosomes of the bacterium Escherichia coli (see Fig. 4) and other bacteria Bacillus subtilis, Rhizobium, etc. (see [27,28]), but they were rarely interpreted as indicating cholesteric packaging of DNA. Data on the state of DNA in germ cells are of interest. In the sperm heads of many mammals (rats, rabbits, stallions, bulls), DNA molecules have cholesteric packaging, despite the presence of protamine-like proteins in their composition. In particular, the circular dichroism spectrum of bull sperm chromatin contains an intense negative band corresponding to the cholesteric packaging of DNA molecules. Similar packaging is characteristic of the sperm of scorpions, octopuses (Eledone cirrhosa), tree frogs (Rhacophoris), and fish (Scyliorhinus caniculis) [27, 28].

In all three cases—in sperm heads, in dinoflagellate chromosomes, and in Escherichia coli—proteins play a significant role in the formation of cholesteric packaging of DNA molecules: protamines (protamines in sperm), histone-like proteins (HLPs), and viral-derived proteins (DVNPs) in dinoflagellates, and Dps in Escherichia coli bacteria. These proteins perform the same function—to compact the DNA. In all cases, the protein reduces the electrostatic repulsion between the negatively charged phosphate groups of DNA, allowing the molecule to fold into a more compact form. This can result in the formation of an ordered, cholesteric liquid crystalline phase.

The Dps protein deserves special mention. The literature generally suggests that the presence of the Dps protein in dormant Escherichia coli cells forms the most densely packed ordered structures—nanocrystalline DNA–Dps complexes [4,16]. It is often assumed that the cholesteric liquid crystalline structure is observed in Escherichia coli only in the absence of DPS in the cell (see Fig. 4A [4]).

#### V.2.2. Model of DNA Structural Organization in Dps-null Cells

To explain the obtained data (Fig. 7), the following structural model of bacterial DNA organization in the dormant state is proposed.

DNA (like Fig. 10A) is organized into parallel sheets (layers), in which long DNA molecules are arranged side by side with a high degree of local parallel ordering. The distance between the centers of adjacent DNA molecules within a single sheet is approximately 44.5 Å. 400 double-stranded DNA fragments with a random nucleotide sequence and a length of approximately 210 bp (≈ 6670 Å) were constructed using the UCSF ChimeraX software package. The fragments were organized into a multilayered supramolecular structure as follows. The structure consists of 20 layers, each containing 20 parallel-oriented DNA fragments. Within a single layer, the fragments were arranged on a lattice with an interaxial distance of 44.5 Å; each fragment was rotated by a random angle around its longitudinal axis. The interlayer distance was 44.5 Å, and the layers were rotated relative to each other by 9° around the plane normal, as in Fig. 10. This geometry ensures a cumulative angular shift in fragment orientation between layers.

The model was visualized using PyMOL. Orthogonal projections of the structure were obtained (frontal, top, top and side views; see Fig. 11 A, B, C). Images of successive virtual sections of the model were also generated. In lateral projections, such sections exhibited characteristic arcuate patterns (Fig. 11 C), caused by the successive rotation of the layers, which is characteristic of the cholesteric packing of DNA molecules. Molecular dynamics simulations were performed in the GROMACS version 2026.0 software package using the AMBER all-atom force field. The system topology was obtained using the pdb2gmx module. The system was placed in an explicit solvent and neutralized with ions. The system energy was minimized using the steepest descent method until a maximum force per atom of no more than 1000 kJ mol⁻¹ nm⁻¹ was reached.

**Figure 11.**
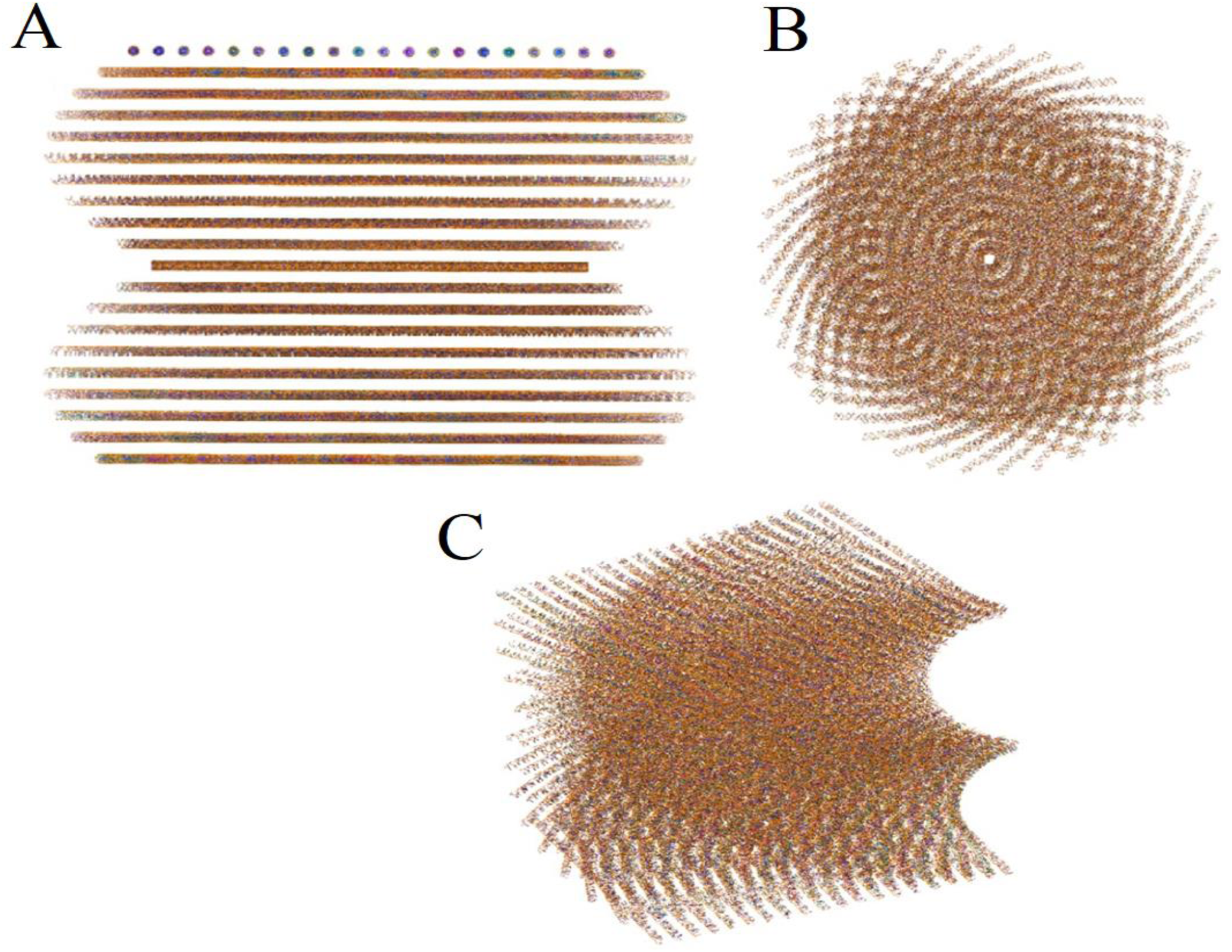
Structural model of the organization of bacterial DNA in a dormant state. The structure consists of 20 layers, each of which contains 20 parallel oriented DNA fragments (similar to Fig. 10A). The distance between the centers of adjacent DNA molecules within a single layer is approximately 44.5 Å. The distance between the layers was 44.5 Å, the layers are rotated relative to each other by 9° around the normal to the plane. A - frontal view of the structure; B - top view; C - top and side view.

The system was then equilibrated. In the first stage (NVT), a v-rescale thermostat (τ = 0.1 ps) was applied for 500 ps at a temperature of 300 K, imposing positional constraints on the DNA atoms. In the second stage (NPT), a Parrinello– Raman barostat (isotropic, τ = 2.0 ps) was used without positional constraints for 500 ps at a temperature of 300 K and a pressure of 1 bar.

The calculation was performed in the NPT ensemble for 4 ns (2,000,000 integration steps with a step size of 2 fs). Electrostatic interactions were calculated using the partial-mesh Ewald (PME) method. Van der Waals interactions were described by a scheme with a cutoff of 1.2 nm. Bonds involving hydrogen atoms were constrained using the LINCS algorithm. The temperature was maintained with a v-rescale thermostat (τ = 0.1 ps), and the pressure was maintained with an isotropic Parrinello–Raman barostat. Trajectories were saved every 100 ps, and energies were saved every 10 ps. The center of mass of the system was removed separately for the DNA group and the water plus ion group. Calculation of X-ray powder diffraction patterns from an atomistic model. After completing the molecular dynamics simulation, the atomic structure of the entire assembly was extracted from the final frame of the trajectory. The structure was trimmed along the boundaries of a rectangular lattice with parameters a = 487.3 Å, b = 797.4 Å, and c = 886.0 Å to include all atoms located within the cell.

To calculate the hkl reflections, a rectangular cell with parameters a = 487.3 Å, b = 797.4 Å, and c = 886.0 Å was selected. These parameters correspond to the model geometry: 19 interaxial intervals of 44.5 Å in the layer plane (19 DNA strands along each direction), and 19 interlayer distances of 44.5 Å along the normal to the layers. This cell represents a representative fragment of a cholesterically twisted structure containing a cumulative director rotation of 171° (compare with Fig. 10). A complete set of hkl reflections was generated in Python (NumPy) over a range of interplanar spacings d from 10 to 100 Å. For each reflection, the intensity I(hkl) = |F(hkl)|² was calculated using kinematic diffraction theory. The structure factor was calculated using the formula:

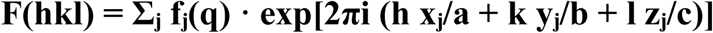

where the summation is over all atoms in the domain, fⱼ(q) is the atomic form factor of the jth atom, which depends on the scattering vector modulus q, and the atomic coordinates are taken directly from the truncated structure. The atomic form factors for the elements H, C, N, O, and P were specified by analytical expressions (Cromer– Mann parameters).

A powder diffraction pattern was constructed based on the calculated intensities I(hkl). To obtain a one-dimensional intensity curve I(q), powder averaging (summing the intensities of reflections with the same scattering vector modulus q) was performed, followed by peak broadening. The broadening was modeled by convolution with a function that considers the finite size of coherent domains of 300 nm (broadening according to the Scherrer equation [26]); and crystal lattice imperfections (paracrystalline disorder and/or microstrains), modeled by an additional Gaussian component. As a result, a realistic one-dimensional scattering curve I(q) was obtained in the q range corresponding to the experimentally observed DNA packing peaks.

The powder diffraction pattern calculated based on this model was directly compared with experimental small-angle X-ray scattering (SAXS) data obtained on Dps-null mutant E. coli cell samples. The comparison result is shown on Fig. 12. It is clear from Fig. 12 that the model proposed in this work completely reproduces the key features of the experimental diffraction pattern from Dps-null mutant E. coli cell samples.

**Figure 12.**
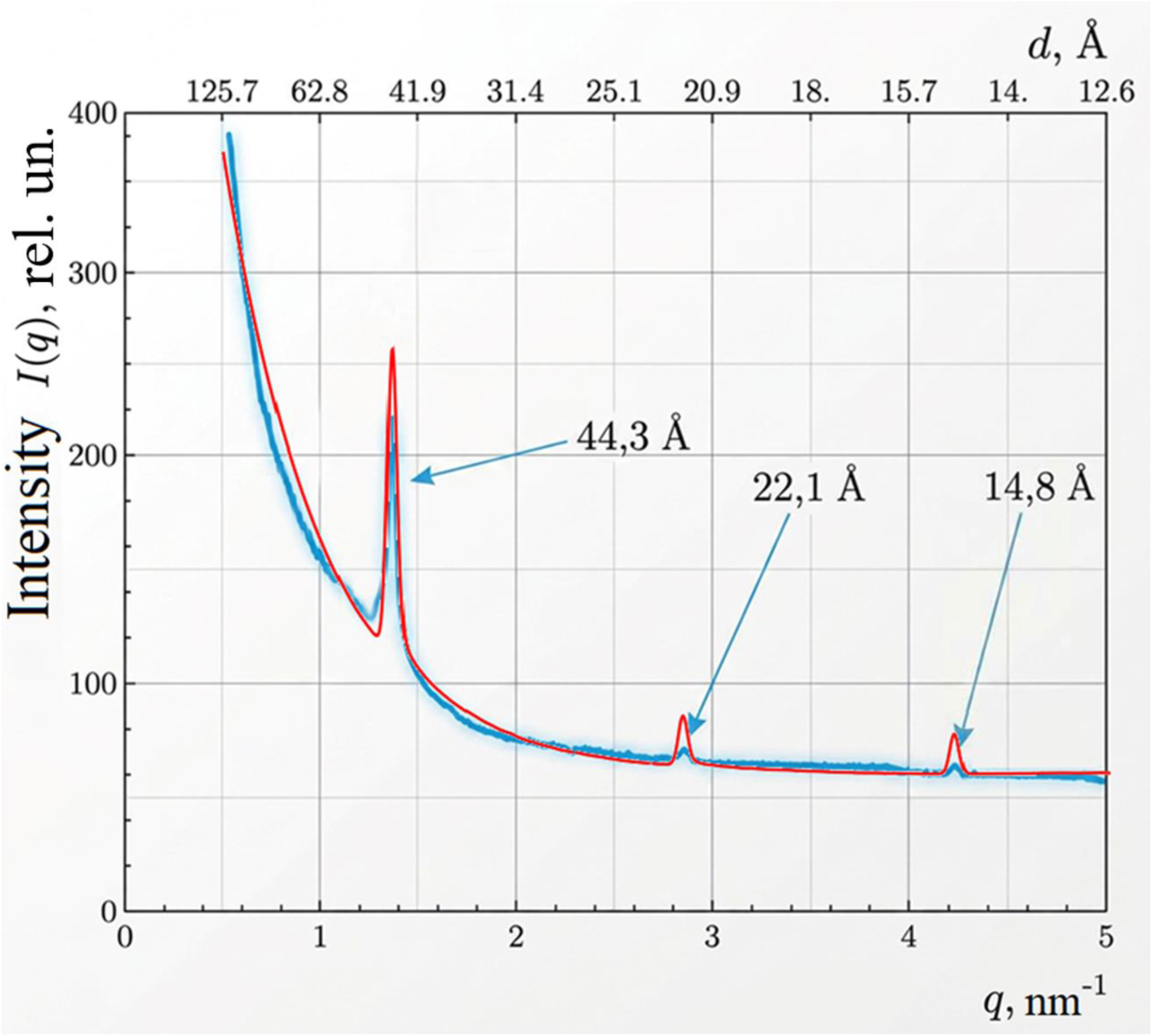
The powder diffraction pattern calculated based on the model (red curve) is compared with the experimental data on the scattering intensity as a function of the scattering vector q (blue curve) for a sample of the Dps-null E. coli strain.

## VI. DISCUSSION

This article presents original experimental results on synchrotron radiation diffraction for samples containing E. coli BL21-Gold (DE3) bacterial cells overproducing the Dps protein, subjected to starvation stress (Fig. 1); upon exposure of E. coli BL21-Gold bacterial cells in the stationary phase to a chemical analogue of the anabiosis autoinducer, 4-hexylresorcinol (4-HR) (Fig. 5); and for samples containing Dps-null K-12 Δdps mutant cells in the stationary phase (Fig. 7). Diffraction maxima are present at ∼44.5, 22.2, and 14.8 Å in all Figs. 1, 5, and 7 (deviations from these values are within the measurement error). These diffraction maxima are also present for samples containing cells of the Dps-null K-12 Δdps mutant, meaning that the Dps protein is not involved in the formation of the diffraction peaks at 44.5, 22.2, and 14.8 Å. These peaks are of greatest interest for understanding DNA packaging and are related to the structural organization of DNA in the cell. The model of the structural organization of DNA as a cholesteric liquid crystal is described in section V.2.2 and describes well the position of the diffraction maxima at ∼44.5, 22.2, and 14.8 Å. Let us turn to the broad maximum at a resolution of 90 Å, shown in Fig. 1 and Fig. 5. If the structural organization of DNA-Dps corresponded to the first model of the appearance of DNA-Dps crystals (Fig. 8a), the distance between the DNA strands would be approximately 90 Å. The diffraction pattern of DNA molecules spaced 90 Å apart should consist of a sharp (narrow and intense) main peak at a resolution of 90 Å and a noticeable satellite corresponding to a resolution of ∼30 Å. A simulation of such a pattern is shown in Fig. 9. Nothing similar was observed in our experiments (see Figs. 1, 5, 7). Therefore, the distance between DNA molecules should be ∼44.5 Å, and the DNA molecules are organized like a cholesteric liquid crystal. Now let’s turn to the Dps protein, which is present in the experiments shown in Fig. 1 and Fig. 5. The Dps protein is, to a rough approximation, a sphere with a diameter of 90 Å. If a crystal were formed during the Dps–Dps interaction (see Fig. 8 A and B), then the diffraction pattern of Dps–Dps, located relative to each other at a distance of 90 Å, should in this case also consist of a main narrow and intense peak with a resolution of 90 Å and a noticeable satellite corresponding to a resolution of 30 Å.

In the experiment at a resolution of 90 Å, a broad peak is observed instead of a narrow and intense one (see Figs. 1, 5). This suggests that the Dps–Dps system, when interacting with DNA, forms a paracrystalline or even amorphously ordered state [26] with the first broad peak of the radial distribution function equal to ∼90 Å. The remaining peaks are strongly smeared (they are wide and low-intensive) [26], therefore they are not visible in the experiment.

Note that the synchrotron radiation diffraction data provides an integrated characteristic of a sample consisting of a collection of individual cells. This means that all types of condensation observed in individual cells and listed in the Table1 are taken into account, namely, nanocrystalline DNA structures, liquid crystalline DNA structures, and folded nucleosome-like DNA structures (see Fig. 3 a, b, and c, respectively). As mentioned earlier, the diffraction pattern of synchrotron radiation scattering from a folded nucleosome-like structure is like the diffraction pattern from a growing cell; that is, it has no diffraction peaks. Therefore, the peaks in the experimental scattering curves (see Figs. 1 and 5) account for contributions only from nanocrystalline and liquid crystalline DNA structures. Interpretation of experimental data on Dps-null K-12 Δdps mutant cells revealed that synchrotron radiation scattering data indicate only cholesteric liquid crystalline DNA packaging in the cell and do not indicate nanocrystalline packaging in the integrated spectrum. The Dps–Dps system bound to DNA contributes to the spectrum as a paracrystalline or amorphously ordered state.

Let us turn to transmission electron microscopy (TEM) data. In [12], three types of new (compared to growing cells) condensed structures were discovered: nanocrystalline DNA structures, liquid crystalline DNA structures, and folded nucleosome-like DNA structures (see Fig. 3 a, b, and c, respectively). Fig. 2 is an example of what is called a nanocrystalline structure in a cell [12]. Figure 2 clearly visualizes the Dps protein, which forms a visible nanocrystalline structure. A filtered DNA–Dps crystal is shown in Fig. 2b. The inset to this figure shows the intensity profile of the electron density along the white line and the densities attributed to the interlayer DNA chains are highlighted [12]. The authors of [16] put forward the hypothesis that DNA is localized between hexagonally packed layers of Dps. From Fig. 2b it is clearly seen that the DNA itself and its structural organization are not visualized; there are only hypotheses that the DNA is localized between hexagonally packed layers of Dps (see Fig. 2b and Fig. 8), following the ideas presented in [16].

Let us turn to data on the liquid crystal structure of DNA. It is noteworthy that liquid crystal structures (illustrated in Fig. 4) were discovered in dormant (starvation stress) E. coli bacterial cells [12] in all their populations: both with and without the dps gene (Dps null), i.e., in the absence of the Dps protein in the cell, although it was previously believed [4,16] that a liquid crystal structure with characteristic arcs and arches can only be observed in the absence of the Dps protein. In some cells, the structure of condensed DNA resembles a cholesteric liquid crystal [4,12]. This is most beautifully illustrated by Fig. 4A, where the characteristic arches are clearly visible.

Fig. 6A.B shows liquid crystal structures for anabiotic dormant cells. These figures also clearly visualize the DNA in the cell. The different radii of the arcs (or arches) observed in Fig. 6A and Fig. 6B indicate different pitches of the cholesteric helix for these structures (or heterogeneity of DNA packaging in different cells). The presence of different amounts of Dps protein in these cells indicates that Dps protein is not an absolute obstacle to the formation of cholesteric liquid crystalline DNA packaging in cells. The cell in Fig. 6B overexpresses Dps, and Dps protein is present in excess.

This statement is also supported by the data in Table 1, which shows that the presence of Dps protein is not an absolute obstacle to the formation of cholesteric liquid crystalline DNA packaging in cells. This also indicates that changes in external conditions and cell strain alter the DNA packaging pattern.

Changes in the environment influence the structural organization of DNA in the cell. It can be assumed, however, that these changes most likely do not affect the hierarchy of DNA structure in the cell nucleoid (see Introduction and [2]). TEM is used to study the structure of DNA in thin 2D cell sections. The structure visible in the section is insufficient to describe the 3D structure of DNA packaging. Therefore, the TEM method used in this study can only visualize structures belonging to the lower (first) hierarchical level of DNA compaction. DNA conformation is not visualized by the TEM methods used in this article in any of the condensed structures except for the liquid crystal.

Let us turn to the experiment performed in [16]. It was shown that at the onset of starvation stress (24 hours), DNA toroids (an intermediate stage of DNA packaging during stress) appear in the cell, and DNA-Dps nanocrystals begin to grow on their surface during further starvation (36, 48 hours). DNA toroids, however, should also appear at the initial stage of starvation in cells lacking the Dps protein. Then, upon further starvation, the intermediate structure—a toroid within which DNA chains form a hexagonal packing [41]—can transform into a cholesteric liquid crystal. This is a possible alternative to the nanocrystalline packaging of DNA-Dps, in which only the crystalline packing of Dps is visible [16]. This alternative requires further clarification using state-of-the-art electron microscopy, X-ray diffraction, and other methods.

In diffraction studies using a synchrotron radiation source, information is read from the entire cell, then averaged over the entire cell ensemble (the entire sample), in another words information is obtained from the entire 3D structural organization of DNA within the cell. In this sense, diffraction studies using a synchrotron radiation source reflect the structural organization of DNA within the cellular 3D structure. Therefore, the observed cholesteric liquid crystalline packaging of DNA reflects its presence throughout the entire 3D volume of the cell.

## VII. CONCLUSIONS

This article presents and critically reviews the results of experimental, original, and literature-based studies conducted by the authors between 2016 and 2026 on the structural organization of DNA in dormant (starvation stress), anabiotic dormant (4-HR treatment) E. coli cells, and the K-12 Δdps strain, which lacks the Dps protein (Dps-null E. coli). The experimental data include small-angle synchrotron radiation diffraction (SAXS) and transmission electron microscopy (TEM) data. It should be noted that SAXS data are obtained under more natural conditions for cell samples than using TEM. Synchrotron radiation diffraction experiments on K-12 Δdps cells (Fig. 7) led to the conclusion that peaks at resolutions of 44.3, 22.1, and 14.8 Å are only associated with an ordered DNA organization. This ordered DNA organization also extends to samples of dormant (starvation stress) cells (Fig. 1) and anabiotially dormant cells (Fig. 5).

The presence of a clear first-order intensity (Fig. 7) and the absence of pronounced higher-order intensity indicates the presence of long-range ordering in only one direction, which is typical for systems with one-dimensional (1D) order and significant disturbances in the other two directions.

This paper proposes a model for the structural organization of bacterial DNA in the dormant state as a cholesteric liquid crystal. The powder diffraction pattern calculated using this model is compared with experimental small-angle X-ray scattering (SAXS) data obtained on Dps-null cell samples. The proposed model fully reproduces the key features of the experimental diffraction pattern from Dps-null cell samples.

Accordingly, the cholesteric liquid crystal model should also apply to DNA packaging in dormant and anabiotically dormant cells. The presence of a broad diffraction peak at a resolution of 90 Å in the spectra (Fig. 1 and Fig. 5) is associated with the presence of the Dps protein. The Dps-Dps system, when interacting with DNA, forms a paracrystalline or even amorphously ordered state with the first broad peak of the radial distribution function equal to ∼90 Å. The remaining peaks are broad and low-intensive, so they are not visible experimentally.

The heterogeneity of DNA packaging is determined by the different pitches of the chiral helix, as seen in Fig. 4 and Fig. 6, since the radii of the archs and arcs in all figures are different.

A characteristic feature of transmission electron microscopy (TEM) data is that DNA is clearly visualized in only two cases: when the DNA is present in the cell in the form of toroids [16] or when the DNA forms liquid crystalline structures (see Fig. 4 and Fig. 6). In the remaining cases, TEM visualizes nanocrystalline structures in the cell formed by the Dps protein. Since DNA is bound to Dps, in vitro experiments on DNA-Dps crystals produce images very similar to the structures shown in Fig. 8 [15, 25], there is a fairly general opinion, which until recently was also held by the authors of this article, that in most cells in the dormant and anabiotically dormant states, the structural organization of DNA is nanocrystalline (see, for example, [4, 7–10, 12–16]). TEM provides a rich picture of the DNA-Dps packaging in short-range order, but says nothing about the long-range order of DNA packaging. The SAXS method does indicate long-range order and indicates the presence of long-range ordering in DNA packaging in only one direction, which is typical for systems with one-dimensional (1D) order.

This article concludes that cholesteric liquid crystalline DNA packaging predominates in dormant and anabiotically dormant cells. The SAXS method does not detect any presence of nanocrystalline DNA packaging in the experiments described here. The Dps protein visualized by TEM may exist in a nanocrystalline state, with the Dps protein obscuring the true DNA conformation. The broad diffraction peak in SAXS associated with the presence of the Dps protein does not confirm the TEM data and suggests that the Dps-Dps system forms a paracrystalline or amorphously ordered state when interacting with DNA.

To determine which structural organization of DNA predominates in the cell—cholesteric liquid crystalline or nanocrystalline—or whether they coexist and fully manifest themselves under different external conditions, recent structural advances should be utilized. These include nanoscale imaging and tomography methods used at the ESRF-EBS synchrotron (Grenoble, France). These methods enable quantitative assessment of the 3D structure and elemental composition of samples in their natural state [42]. In transmission electron microscopy, an improved method for DNA detection using a fluorescent dye (ChromEMT) is used to visualize chromatin in situ [43].

The model proposed in this work is a useful and effective tool for interpreting experimental diffraction data and allows quantitatively linking the atomistic organization of DNA with the macroscopic diffraction features of the liquid crystalline state in a living cell.

## ACKNOWLEDGMENTS

X-ray diffraction measurements were performed using synchrotron radiation at the ESRF beamline ID23-1 (Grenoble, France). The authors are grateful to the ESRF for providing the opportunity to conduct the experiments.

Some repeat measurements were conducted at the BioMUR (BioSAXS) station of the Kurchatov Synchrotron and Neutron Research Complex. The authors are grateful to the Kurchatov Synchrotron Research Center for the opportunity to conduct the experiments.

Analytical electron microscopy and biaxial tomography were performed at the “Electron Microscopy in Life Sciences” User Center at Lomonosov Moscow State University.

## DECLARATIONS

### COMPLIANCE WITH ETHICAL STANDARDS

This work does not involve human or animal studies.

